# DNA Topoisomerase I differentially modulates R-loops across the human genome

**DOI:** 10.1101/257303

**Authors:** Stefano Giustino Manzo, Stella Regina Hartono, Lionel A. Sanz, Sara De Biasi, Andrea Cossarizza, Giovanni Capranico, Frederic Chedin

**Affiliations:** Department of Pharmacy and Biotechnology, University of Bologna, Bologna, Italy; Department of Molecular and Cellular Biology and Genome Center, University of California, Davis, United States; Department of Medical and Surgical Sciences for Children and Adults, University of Modena and Reggio Emilia, Modena, Italy

## Abstract

**Background:** Co-transcriptional R-loops are abundant non-B DNA structures in mammalian genomes. DNA Topoisomerase I (Top1) is often thought to regulate R-loop formation owing to its ability to resolve both positive and negative supercoils. How Top1 regulates R-loop structures at a global level is unknown.

**Results:** Here, we performed high-resolution strand-specific R-loop mapping in human cells depleted for Top 1 and found that Top1 depletion resulted in both R-loop gains and losses at thousands of transcribed loci, delineating two distinct gene classes. R-loop gains were characteristic for long, highly transcribed, genes located in gene-poor regions anchored to Lamin B1 domains and in proximity to H3K9me3-marked heterochromatic patches. R-loop losses, by contrast, occurred in gene-rich regions overlapping H3K27me3-marked active replication initiation regions. Interestingly, Top1 depletion coincided with a block of the cell cycle in G0/G1 phase and a trend towards replication delay.

**Conclusions:** Our findings reveal new properties of Top1 in regulating R-loop homeostasis and suggest a potential role for Top1 in controlling replication initiation via R-loop formation.

## Background

Biological processes such as transcription and replication generate torsional stress on the DNA double helix that, if not properly dealt with, can lead to genome instability [1]. R-loop structures, aprevalent non-B DNA structure in mammalian genomes, have been particularly linked to genomic instability by causing interference between the replication and transcription machineries [2, 3]. R-loops are formed during transcription upon reannealing of the nascent transcript to the DNA template strand, forming an RNA:DNA hybrid and forcing the non-template strand to loop out. Mapping data indicate that these non-B DNA structures are prevalent in mammalian genomes, where they form dynamically over conserved regions [4, 5]. Negative supercoiling generated behind the elongating RNA polymerase [6] is thought to facilitate R-loop formation by inducing an underwound DNA state favorable to the re-annealing of the nascent transcript [7]. DNA Topoisomerase I (Top1) is one main cellular factor controlling topological homeostasis [8, 9]. Top1 activity can relax negative supercoils by cutting one of the DNA strands, creating a transient Top1-DNA cleavage complex (Top1cc), and performing a controlled rotation of the cut strand around the uncut strand [10, 11]. The relaxation activity on negative supercoils is thought to reduce co-transcriptional R-loop formation which in turns prevents replication / transcription interference and favors genome stability. Indeed, deletion of the bacterial *topA* gene, an enzyme that only relaxes negative supercoils, creates R-loop-prone hypernegatively supercoiled DNA and causes a growth defect that can be suppressed by over-expression of Ribonuclease H (RNase H), an enzyme that degrades RNA strands in RNA:DNA hybrids [7, 12, 13]. Furthermore, persistent depletion of Top1 in mammalian cells leads to replicative stress and replication-transcription conflicts that can be rescued by overexpression of RNase H [14]. Finally, stabilization of Top1cc by Top1 inhibitors such as camptothecin and its derivatives [15] leads to R-loop stabilization in human cells upon short treatment [16, 17] and to transcription-dependent DNA breakage that can be partially suppressed by RNase H expression [18].

Thus, while it is clear that Top1 regulates R-loops and prevents R-loop-induced genomic instability, the range of loci that are sensitive to R-loop modulation by Top1 is not known. Addressing this gap in knowledge is important given rising evidence that R-loops are abundant in mammalian genomes and also participate in important biological processes [19-21]. For instance, R-loops are involved in regulating chromatin states [4, 5, 22], in mediating transcription termination [23], and in immunoglobulin class switch recombination [24]. Studies also suggest a role for R-loops in priming DNA replication in prokaryotic systems and yeast [25-28]. How R-loop formation is dynamically regulated to permit the physiological roles of R-loops while minimizing the negative impacts of excessive R-loops on genome stability is not clear. In this study, we used the DRIPc-seq technique [4] to map R-loop structures genome-wide in human cells experiencing an acute but transient depletion of Top1. Our work reveals that Top1 modulates R-loop structures differently according to genomic context and provide new evidence that R-loops can play a role in the replication process.

## RESULTS

### Top1 depletion causes subtle R-loop gains

To investigate how global R-loop patterns change upon Top1 depletion, we transiently silenced Top1 in human HEK293 cells down to undetectable levels (Fig. 1A) and quantified the global levels of R-loops using dot blots, taking advantage of the anti RNA/DNA hybrids S9.6 antibody [29, 30]. This approach reproducibly showed a 1.5-2-fold increase in overall R-loop loads in Top1-depleted cells (Fig. 1B and Supplemental Fig. S1A). As a control, we pre-treated the genomic DNA with RNase H, which completely abolished the R-loop signal in both control and Top1 knockdown samples, demonstrating specificity of these methods (Supplemental Fig. S1B).

**Figure 1.**
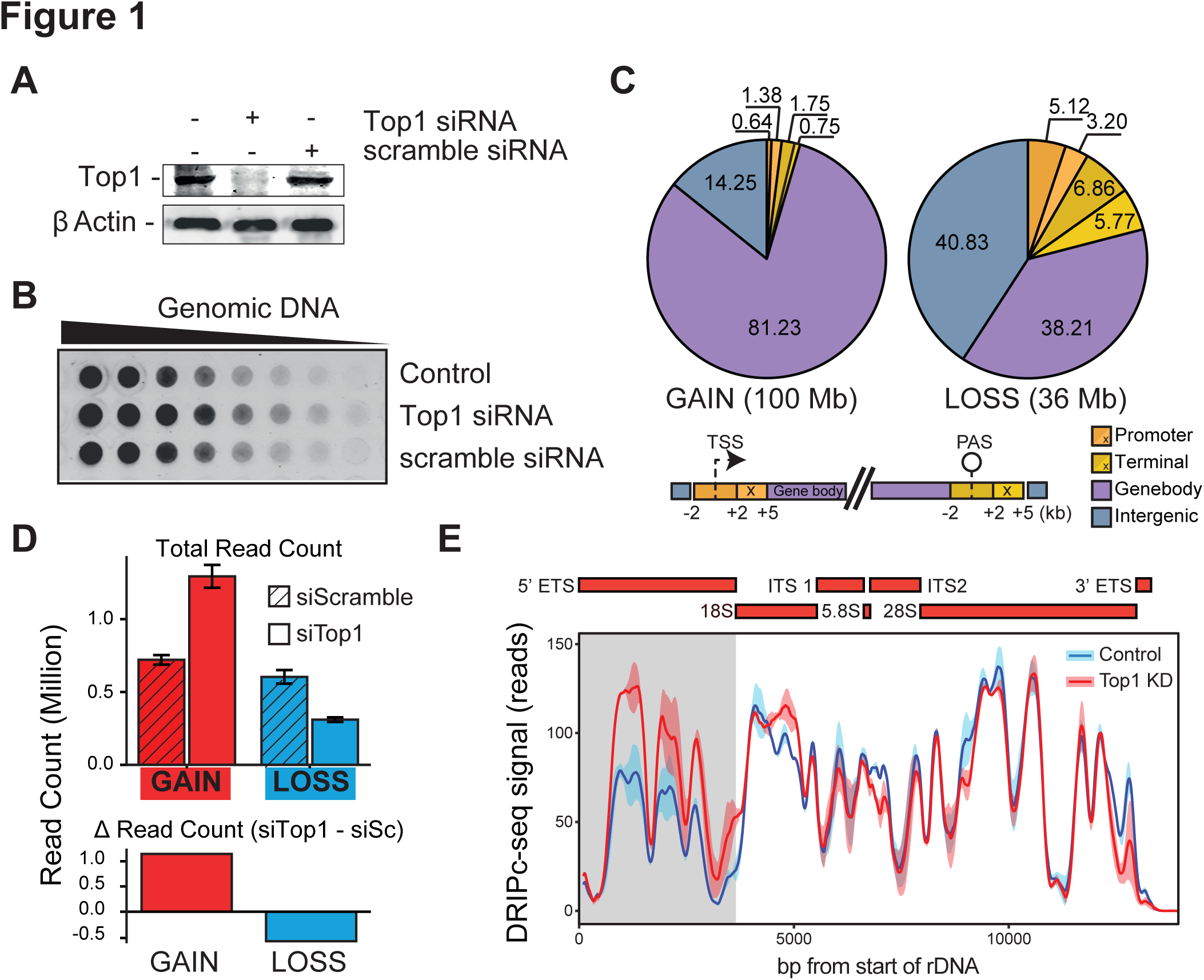
Topoisomerase 1 depletion instigates R-loop changes genome wide. (A) Western blot verifying Top1 depletion upon specific siRNA transfection compared to control β-actin. (B) Dot blot analysis of R-loop formation: two-fold serial dilutions of genomic DNA were arrayed on a membrane and probed using the S9.6 antibody. (C) Distribution of DRIPc peaks gains (left) and loss (right) upon Top1 depletion across several genomic compartments depicted below. Numbers indicate the percentage occupied by each compartment. The total genomic space covered by R-loop gains and losses is indicated. (TSS) transcription start site; (PAS) poly-adenylation site. (D) Top; total number of uniquely mapped reads overlapping with peaks of R-loop gains (left) and losses (right) in control and Top1-depleted samples. Bottom, the relative difference in reads between gains and losses indicates that gains predominate over losses. (E) DRIPc-seq signal profiles for control and Top1-depleted cells over rDNA region. Average signal over two replicates is shown as solid line with standard error (shaded). Structural features of the rDNA region are on top. The 5’ETS region shows significant R-loop increase (grey shade).

We next quantified R-loop formation genome wide by employing DRIPc-seq, a technique that allows high-resolution, strand-specific genomic mapping of R-loop structures [4]. R-loop formation was prevalent in HEK293 cells, giving rise to ~70,000 peaks and covering ~200 megabases (Mb) of genomic space, which is in close agreement with previous data [4]. As expected, R-loop formation was predominantly genic, with promoters and terminators representing hotspots of signal (Supplemental Fig. S1C). Detection of R-loop signal changes after standard peak calling [4] indicated that only a small subset of R-loop peaks (4%) showed significant changes upon Top1 depletion. However, inspection of signal in Top1-depleted samples revealed numerous instances of signal spreading from existing R-loop peaks. Since such events were not properly called, we optimized a higher sensitivity version of our peak-calling algorithm (see Methods). Using this method, we identified significant and reproducible R-loop signal gains and losses upon Top1 depletion, occurring at over 28,000 peaks, or 7% of total peaks (Supplemental Fig. S1D, Supplemental Table 1). These changes were independently validated using DRIP-qPCR at representative test loci (Supplemental Fig. S1E). Similar results were obtained when we induced Top1 depletion with an additional siRNA (Supplemental Fig. S1F). Quantitative analysis of DRIPc-seq signal at these R-loop gain (RLG) and R-loop loss (RLL) peaks was consistent with the increased S9.6 signal detected by dot blot: RLG peaks occupied ~100 Mb of sequence space whereas, RLL peaks occupied 36 Mb only (Fig. 1C). Furthermore, the intensity of R-loop signals measured over all RLG and RLL peaks also showed an overall net increase in Top1-depleted cells (Fig. 1D). This increase in genic R-loops was further augmented by RLG over the 5’ ETS, but not the transcribed 28S region of the ribosomal DNA (Fig. 1E, Supplemental Fig. S1G). Similar results were obtained with a second siRNA for Top1 (Supplemental Fig. S1H). Ribosomal DNA arrays harbor a major source of cellular R-loops [19] and Top1 depletion in yeast was previously shown to cause R-loop gains over the 5’ETS region [31]. Therefore overall, Top1 depletion results in a net increase in cellular R-loop loads, although only a minority of R-loop peaks are directly affected.

### R-loop gains and losses in Top1-depleted cells define distinct gene categories

Peaks of RLG and RLL in Top1-depleted cells appeared to have distinct genomic distributions. RLG peaks overwhelmingly (>80%) matched to gene body (Fig. 1C), indicating that they associate with transcription elongation. RLL peaks, by contrast, matched to promoter and terminator regions, with only a minority (38%) mapping to gene bodies. Moreover, RLL peaks most often showed an “intergenic” distribution that often corresponded to a terminator-downstream location immediately outside of the gene boundaries used here for classification. Mapping RLG and RLL peaks onto genes identified RLG genes that predominantly gained R-loops, as well as RLL genes that mostly lost R-loop signal (Supplemental Fig. S2A and Supplemental Table 1). In addition, a large class of genes showed mixed gains and losses (Supplemental Fig. S2B and Supplemental Table 1). To simplify the analysis, we selected RLG genes with a minimal 5:1 ratio of peak gains to loss (and vice-versa for RLL genes). This delineated two clear groups of RLG (n=959) and RLL genes (n=2046), respectively (Fig. 2A). A large fraction of genes underwent mixed changes (gain and loss; n=9,375), while a small portion did not undergo detectable changes (n=2,464) (Fig. 2A). Altogether, Top1 depletion triggered subtle shifts in genic R-loop distribution marked by both gains and losses of R-loops over a large portion of genes. Overall, a significant subset of genes (11.8% of total) was marked by nearly exclusive patterns of R-loop gains or losses upon Top1 depletion.

**Figure 2.**
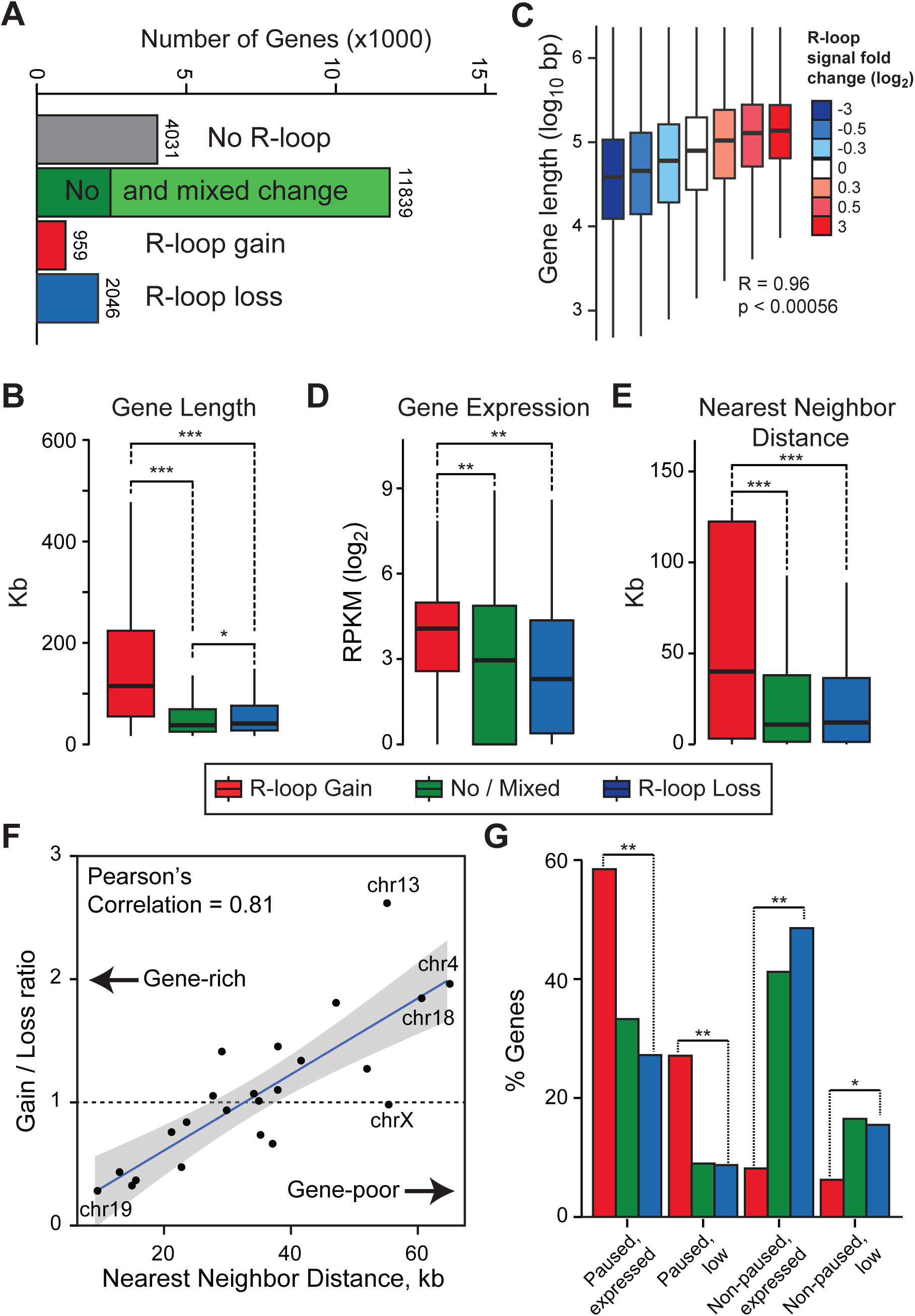
R-loop gains and losses occur on genes with distinct categories. (A) Breakdown of genes according to their R-loop status upon Top1 depletion; numbers indicate gene numbers in each category. (B) Quartile plot depicting distribution of lengths for genes undergoing R-loop gains, losses, or no/mixed change (color code is indicated below the plot). Stars (*, **, and ***) indicate p-value less than 10-^10^, 10-^25^,10-^40^, respectively (Wilcoxon Mann-Whitney). (C) Gene length is plotted as a function of the fold R-loop signal change. Bins were chosen so they contain similar number of genes to avoid biased sampling. Median Pearson correlation coefficient and associated p-value are indicated. (D, E) Same as (B) except gene expression and gene distance are plotted. (F) XY plot between gene density measured on each individual chromosome (represented by a dot) and the ratio of gene numbers undergoing R-loop gains and R-loop losses upon Top1 depletion. The regression line along with 95% confidence interval and Pearson correlation coefficient are indicated. (G) Distribution of genes undergoing R-loop gains and losses according to the RNAP stalling and expression status of each gene. Color code is as in (B).

To understand the nature of the differential response to Top1 depletion, we focused on RLG and RLL genes and first investigated their lengths. RLG genes were significantly longer (2.7 fold on average) compared to RLL genes or genes with no or mixed R-loop change (Fig. 2B). By contrast, RLL genes were only slightly longer than control genes with no or mixed changes in R-loops. This suggests that longer genes are more prone to RLG upon Top1 depletion. To confirm this, we clustered all R-loop peaks by their ratio of R-loop signal change from control to Top1-depleted conditions regardless of whether they belonged to RLL or RLG genes. We then measured gene lengths as a function of R-loop signal fold change in each cluster. We observed a strong positive correlation between the relative intensity of R-loop signal changes and each cluster’s length (Fig. 2C). This, together with the marked distribution of RLG peaks to gene bodies, suggests that transcription elongation through long genes is more prone to R-loop stabilization in absence of Top1. To determine if expression levels could also distinguish RLG and RLL genes, we performed total RNA-seq on both control and Top1-depleted cells. This analysis revealed that RLG genes were significantly more expressed (1.5-3.5 fold on average) than RLL genes, which themselves were expressed on par with control genes (Fig. 2D). Thus, RLG genes tend to be long and highly expressed. Since long genes often reside in gene-poor areas of the genome, we measured the distance between RLG and RLL genes and their nearest neighbor. RLG genes were located significantly further away from potential neighbors than control or RLL genes (Fig. 2E). This indicates that RLG genes tend to occupy gene-poor neighborhoods. Consistent with this, there was a strong inverse correlation between chromosomal gene density and R-loop loss/gain ratio: gene-rich chromosomes predominantly showed loss of R-loop signal loss following Top1 knockdown, while gene-poor chromosomes favored R-loop gain events (Fig. 2F). Finally, given that Top1 can regulate RNAP release from promoter-proximal pausing [32-34], we asked whether RLG and RLL genes differ in how frequently they undergo pause-release. For this, we performed RNA Polymerase II ChIP-seq in control and Top1-depleted cells and determined the pausing index of each gene according to well-defined categories [35]. RLG genes were strongly enriched in genes that undergo promoter-proximal pausing (Fig. 2G). By contrast, RLL genes mostly corresponded to genes that do not undergo pausing and their distribution was not significantly different from that of control genes. More broadly, our analysis confirmed that Top1 depletion causes RNAP accumulation in the vicinity of the TSS, particularly for paused genes [32] (Supplemental Fig. S2C). Altogether, this shows that the differential response to Top1 depletion defines two broadly distinct classes of genes.

### Top1 depletion favors co-transcriptional R-loop gains

The preferential localization of RLG peaks to gene bodies suggests that R-loop gains occur during transcription elongation. Gene metaplots confirmed that RLG peaks were typically circumscribed to the transcribed portion of long genes, with no apparent gradient towards the 5’ or 3’-ends (Fig. 3A). By contrast, RLL peaks were more prominent for shorter genes and showed preferential distribution around the promoter and terminator regions. To determine if the increased R-loops over RLG genes was caused through a co-transcriptional mechanism, we asked if the strandedness of RLG peaks was concordant with the strandedness of their respective genes. Gains of R-loop signal were indeed only observed on the template strand (Fig. 3B). By contrast, the patterns of R-loop signal loss over RLL genes were more complex, even in control cells (Fig. 3C). This complexity most likely reflects the fact that RLL genes reside in gene-rich areas and therefore often possess immediate neighbors with divergent promoters and convergent terminators. As a result, R-loop formation appears on both template and non-template strands in metagene plots. Under conditions of Top1 depletion, both template and non-template signals were reduced. In agreement with their lower transcription levels, RLL genes also showed a lower median R-loop signal compared to RLG genes, (compare Fig. 3B and 3C). Gene expression analysis shows that the large majority (93%) of RLG and RLL genes do not undergo significant up or downregulation in their expression levels, arguing that the changes described here are not simply a result of altered transcriptional states (Supplemental Fig. S2D). Overall, the data are consistent with R-loop changes triggered by Top1 depletion being mainly co-transcriptional.

**Figure 3.**
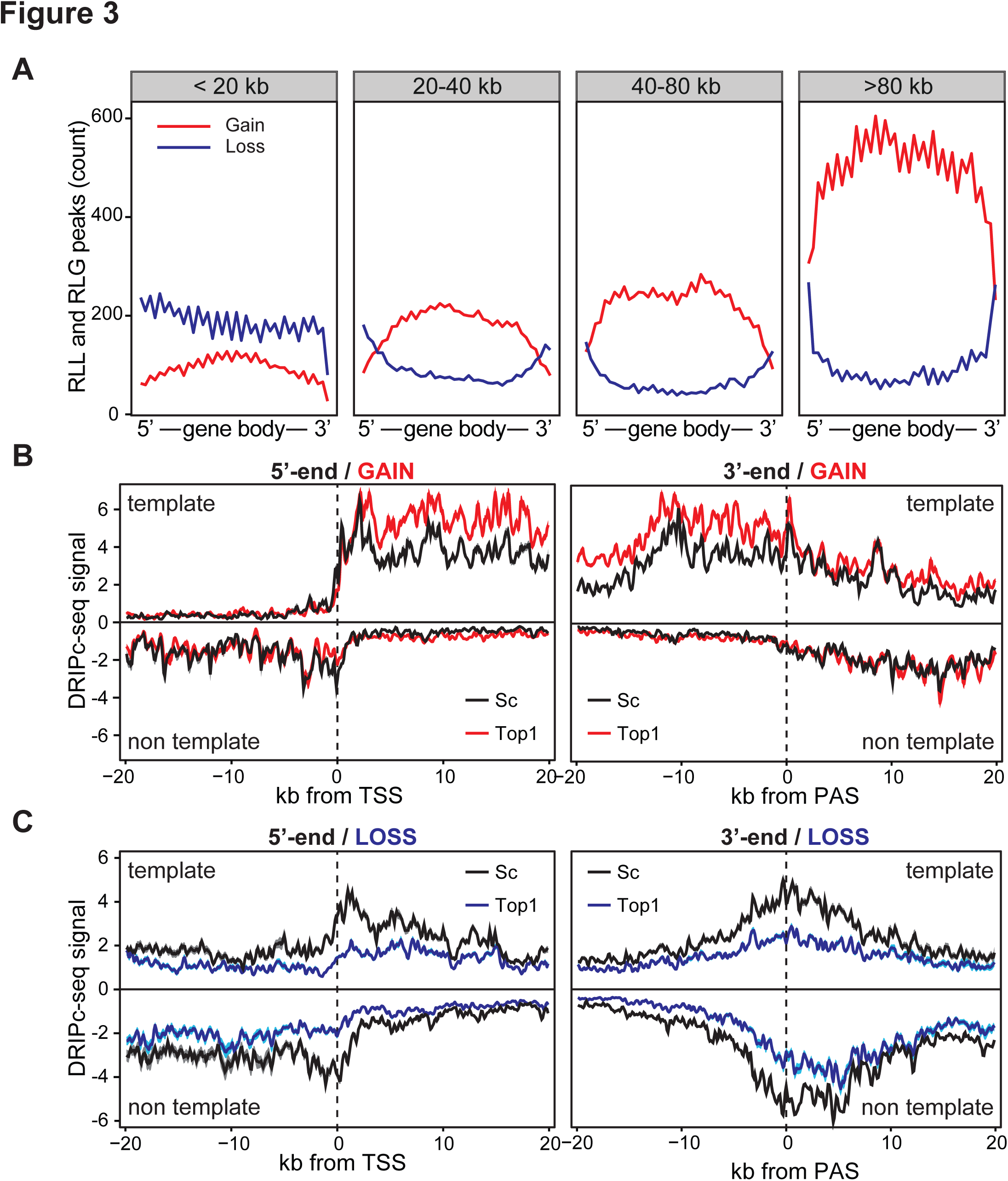
R-loop signal at RLG genes are co-transcriptional. (A) Distribution of all peaks of R-loop gain and loss (p-adjusted < 0.1) along a gene metaplot. Genes are broken down by length, as indicated at top. Genes were binned in 40 bins and peak counts reported by bin. (B-C) Metaplots of DRIPc-seq signal over RLG genes (B) and RLL genes (C) along a 20 kilobase window centered on their TSS at left, or PAS at right. Values are median and shown with standard deviation (shaded). Samples are color-coded as indicated.

### R-loop gains upon Top1 depletion preferentially associate with heterochromatin and nuclear lamina

To understand the cause(s) driving RLG in the absence of Top1, we asked if RLG peaks associate with specific chromatin features compared to R-loop-forming loci matched for position, expression, and length that showed no or mixed changes in response to Top1 depletion (thereafter referred to as “matched”). Matching on GC skew and overall R-loop levels was also performed and did not change the conclusions (data not shown). Using this stringent comparison approach, we first analyzed chromatin association by the extent of peak overlap, keeping promoters, gene bodies, and terminal regions separate. Out of a wide array of available datasets including ChromHMM states [36] and ChIP-seq information for nearly a hundred types of histone modifications and chromatin factors [37], RLG peaks showed modest but significant overlap enrichment only for a few chromatin marks, highlighting their significance. H3K36me3 and H3K79me2, two marks associated with transcription elongation, and the ChromHMM transcription elongation state itself, were enriched over RLG peaks (Fig. 4A). RLL peaks, by contrast, were significantly depleted for these marks compared to matched sets. In addition, RLG peaks showed significantly higher overlap with the heterochromatin mark H3K9me3 and with lamina-associated domains (LADs) [38] while RLL genes showed the opposite trend (Fig. 4A). These observations are consistent with RLG genes being long and highly expressed and occupying gene-poor neighborhoods (Fig. 2) that are more likely to associate with heterochromatic regions. Since lamin and H3K9me3 association might represent physical constraints to the dissipation of transcription-induced supercoils in the absence of Top1, we investigated the relationship between RLG and these parameters further.

**Figure 4.**
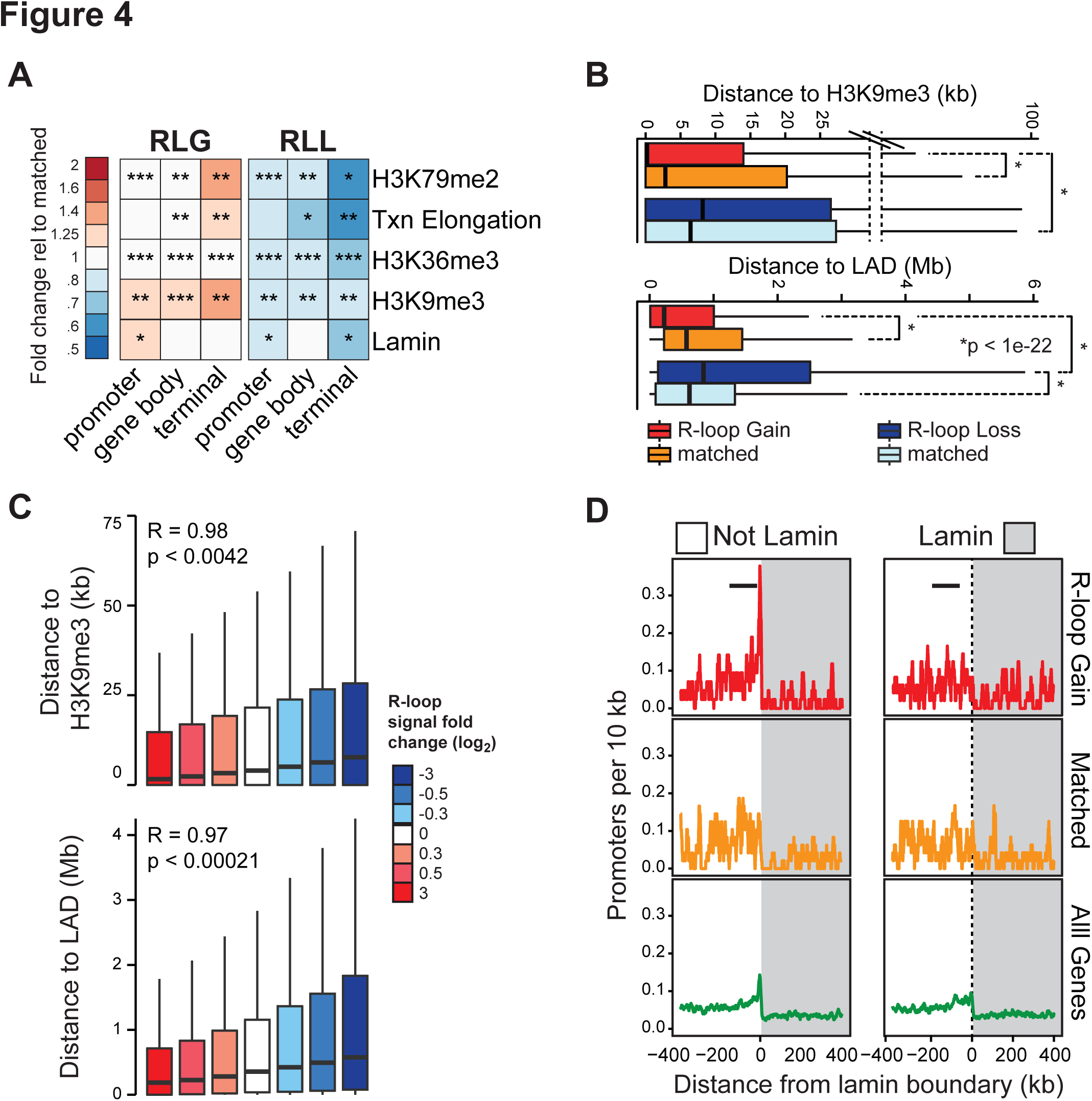
RLG and RLL peaks show distinct epigenetic features. (A) Heatmap indicating the relative enrichment or depletion of RLG and RLL peaks over specific chromatin features shown at right. The ratio of observed over expected overlaps between RLG and RLL peaks and matched R-loop control peaks was measured over each chromatin feature (see Methods) and shown as a color-coded heatmap (shown at left). Stars indicate the extent of overlap between R-loop peaks and each chromatin feature (* 10-25%; ** 25-50%; *** >50%; no star <10%). All values are significant with p-value < 0.008 (Monte-Carlo). (B) Distance between RLG and RLL peaks and H3K9me3 peaks (top) or LADs (bottom) compared to matched controls. Statistical significance was measured by Wilcoxon test. (C) Distance between all R-loop peaks and H3K9me3 peaks (top) or LADs (bottom) after clustering R-loop peaks according to the strength of signal change upon Top1 depletion (color-coded as in Fig. 2C). (D) Promoter density plotted along a region centered on LAD boundaries (shaded) for promoters driving transcription away from the boundary (left) or towards it (right), as indicated by the arrow. Genes were broken down between RLG genes (top), control matched genes (middle) and all genes (bottom).

We calculated the distance between RLG and RLL peaks and the nearest annotated H3K9me3 or LAD peak. RLG peaks were significantly closer to H3K9me3 peaks compared to matched R-loop forming peaks (Fig. 4B). This proximity was true regardless of the genic location of the RLG peak (promoter, gene body and terminator) and was striking given that the median value for distance was close to zero. By contrast, RLL peaks were located further away from H3K9me3 peaks (median distance of 8 kb) than matched control peaks. RLG peaks therefore tend to reside in immediate proximity to H3K9me3 peaks. We next asked if the intensity of R-loop signal gains was correlated to the distance to H3K9me3 peaks. For this, we clustered all R-loop peaks by their R-loop change ratios and measured the distance between these loci and the nearest H3K9me3 peak in each cluster. A strong correlation between the two parameters was observed such that the R-loop peaks with the strongest gains were located closest to H3K9me3 peaks. Vice-versa, R-loop peaks with the strongest losses were located furthest away from H3K9me3 peaks (Fig. 4C). Furthermore in 70% of cases, the H3K9me3 peaks were located upstream of the RLG peaks relative to gene transcription (not shown). Similar observations were made with LADs: RLG peaks were located significantly closer to LADs than RLL genes or control matched genes (Fig. 4B). Similarly, we observed a strong correlation between distance to LADs and strength of R-loop change (Fig. 4C). While the median distance between RLG and annotated LADs was comparatively large (~200 kb; Fig. 4B), we note that top RLG genes in terms of signal gains were often closely juxtaposed to LADs (Supplemental Fig. S3A). Similarly, promoters of RLG genes were characterized by a strong upstream Lamin B1 signal and marked lamin B1 depletion around the TSS region (Supplemental Fig. S3B). To further characterize the arrangement of RLG genes relative to LAD boundaries, we calculated the density of promoters around LAD boundaries as a function of genic orientation. RLG genes transcribing away from LADs showed a sharp promoter density peak at or near the LAD border (Fig. 4D). By contrast, no peak was observed for genes transcribing towards the LAD. A corresponding promoter peak was not observed for matched control genes and only weakly for all genes. Altogether, this data shows that long, highly transcribed genes accumulate R-loops in proximity to H3K9me3 peaks and LADs in the absence of Top1.

### R-loop losses upon Top1 depletion preferentially associate with early, active, replication origins

To gain insights into the functional significance of R-loop loss in the absence of Top1, we analyzed the chromatin states of RLL peaks. Compared to matched R-loop-forming loci, RLL peaks showed significantly higher intersect with chromatin features characteristic of regions repressed by the Polycomb group complex. These features include the “repressed” ChromHMM state, the Polycomb mark H3K27me3, and several subunits of the PRC1 and PRC2 complexes (Fig. 5A). In all cases, a significant decreased overlap was observed for RLG peaks, suggesting that these chromatin features are specific. Additional overlap enrichment was observed for terms related to the chromHMM “insulator” state and for the CTCF and RAD21 factors that often colocalize at chromatin loops (Supplemental Fig. S4A).

**Figure 5.**
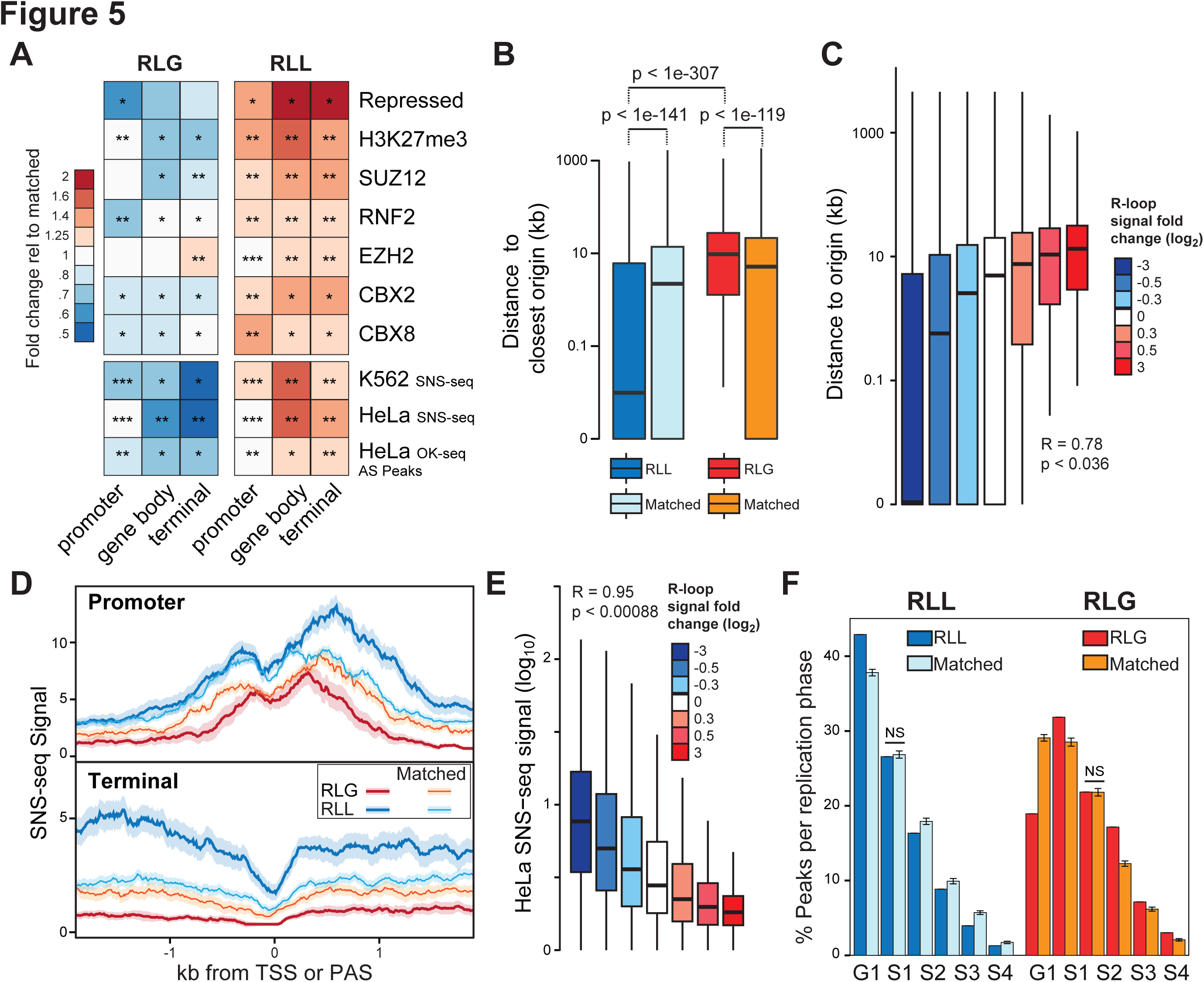
RLL peaks are enriched for replication origins while RLG peaks are depleted. (A) Heatmap of enrichment or depletion of RLG and RLL peaks over specific chromatin features. Color codes and description are as Fig. 4A. (B) Distance between RLG and RLL peaks and replication origins compared to matched controls. Statistical significance was measured by Wilcoxon test. (C) Distance between replication origins and all R-loop peaks ranked by the strength of R-loop changes (color-coded). (D) SNS-seq signal plotted over promoter and terminator regions for RLL and RLG loci as well as matched controls. Data is shown as median with standard error (shaded). (E) SNS-seq replication signal of R-loop peaks ranked by the strength of R-loop gains and losses (color-coded). (F) Replication timing analysis for RLL, RLG and matched peaks according to phases of the cell cycle based on Repli-seq data. All comparisons to matched peaks are significant (p < 0.008, Monte-Carlo) except when indicated (NS).

Since Top1 activity has been mapped to a well-established, conserved human replication origin [39], we also included replication initiation sites mapped by Short Nascent Strand sequencing (SNS-seq) [40] to our analysis. To our surprise, RLL peaks showed strong overlap enrichment with such loci compared to matched controls while RLG peaks showed clear depletion (Fig. 5A). To investigate the relationship between RLL peaks and origins further, we measured the distance between RLL and RLG loci to the nearest annotated SNS-seq loci. RLL peaks were located significantly closer to origins than matched controls and dramatically closer than RLG peaks which themselves were located further away from origins than matched controls (Fig. 5B). Furthermore, we observed a strong correlation between the intensity of the R-loop signal gain/loss upon Top1 depletion measured over RLL and RLG peaks and the distance from these peaks to the nearest SNS-seq origin (Fig. 5C). Strikingly, peaks with the strongest loss tended to directly match onto SNS-seq origins (median distance of zero). Increased overlap of RLL peaks over CpG island loci, which are often replication origins [40], was also observed, while RLG peaks showed the opposite trend (Supplemental Fig. S4A). Thus, topoisomerase I depletion associates with a loss of R-loop signal at peaks that are proximal to replication origins. We next asked if the relationship between RLL peaks and origins also applied to SNS-seq signal, which reflects the frequency of replication initiation events. SNS-seq signal was significantly higher for RLL peaks over both promoters and terminators compared to the signal observed over matched controls (Fig. 5D). Similarly, SNS-seq signal over RLG peaks was lower than that of matched controls. Overall, a strong correlation was observed between replication signal and the strength of R-loop gains and losses (Fig. 5E). This indicates that RLL peaks delineate regions with high replication initiation activity while RLG peaks match further away from sites of replication initiation onto regions with poor replication initiation potential. Analysis of replication timing data (Repli-seq) confirmed that RLL peaks replicate early, mostly in G1 and S1 phases (Fig. 5F). Compared to matched loci, RLL peaks were significantly more likely to replicate in G1 and less likely to replicate in later phases. By contrast, RLG peaks replicated predominantly in later phases of the cell cycle (S1 and S2) and showed a significant tendency towards later replication compared to matched loci. To ensure that the association between RLL peaks and replication origins is robust, we analyzed independent datasets where origins were mapped through Okazaki fragment sequencing (OK-seq) [41]. While SNS-seq and OK-seq datasets produce distinct replication initiation maps, both methods nonetheless show significant overlap in particular around gene bodies and terminal genic regions (data not shown). RLL peaks showed increased overlap with OK-seq-derived initiation peaks (AS peaks, [41]) while RLG peaks showed decreased overlap (Fig. 5A). Likewise RLG peaks were located further away than expected from matched control genes, while RLL peaks were distributed as expected from, or closer than, control peaks (Supplemental Fig. S4B). Finally, RLG peaks showed reduced densities of AS peaks compared to matched controls while RLL peaks showed the opposite trend (Supplemental Fig. S4C). Thus, the findings show a robust association between peaks of R-loop loss in response to Top1 depletion and early replication origins. While the analysis above was restricted to genic regions so we could ensure stringent matching procedures, we determined that intergenic RLL peaks also showed a 4-5 times higher overlap with replication initiation regions (SNS-seq or OK-seq) than expected at random (data not shown).

### Top1 depletion triggers G0/G1 block and replication timing delays with minimal DNA damage

Since R-loop accumulation, as seen here over long genes in Top1-depleted cells, often associates with increased genomic instability, we tested if Top1 depletion was associated with induction of the DNA damage response. Surprisingly, Western blots did not indicate significant hyper-phosphorylation of the histone variant H2AX, a marker of DNA breaks and replicative stress (Fig. 6A). Camptothecin treatment, which leads to covalent Top1-DNA complexes, strongly induced this modification (Fig. 6A). A more sensitive examination of γH2AX levels by immunocytochemistry revealed a small but significant increase in Top1-depleted cells (Fig. 6B). However, no significant increased phosphorylation of ATM, CHK1, or CHK2 could be measured by Western blots, suggesting that ATM and ATR DNA damage sensing pathways are not broadly activated upon acute Top1 depletion in human HEK293 cells (Fig. 6A). Since DNA damage induced by Top1 depletion is dependent on S-phase [14], we next asked if Top1 depletion caused any cell cycle changes by performing cytofluorimetric analyses. Surprisingly, Top1 depletion induced a consistent block in G0/G1 phase, with marked reduction of the S and G2/M phases (Fig. 6C), which was not due to checkpoint activation (Fig. 6A). Immunofluorescence assays using the Ki-67 proliferation marker confirmed that a significant fraction of Top1-depleted cells exit out of the cell cycle and rest in G0 (Fig. 6D and E). Similar results were obtained using an independent Top1 siRNA (Supplemental Fig. S5). Thus, a pronounced, short term (5 days), depletion of Top1 impairs the G1 to S phase transition. This observation may account for the modest amount of DNA damage observed under these conditions since R-loop mediated genomic instability is thought to be caused by replication-transcription conflicts. Indeed, prolonged Top1 depletion under selective conditions that force cells to undergo division is associated with R-loop-induced DNA damage [14]. Given the accumulation of Top1-depleted cells in G0/G1, we wondered if the R-loop gains and losses we observed could be due to preferential R-loop formation by RLG and RLL genes within and outside of G1, respectively. To test this, we synchronized cells in G2, released them, and monitored R-loop formation in G1 and mid-S by DRIP-qPCR at a range of loci. Other than an expected drop of R-loop formation in G2/M, we did not observe any specific trend across RLG and RLL loci analyzed here (Supplemental Fig. S6A).

**Figure 6.**
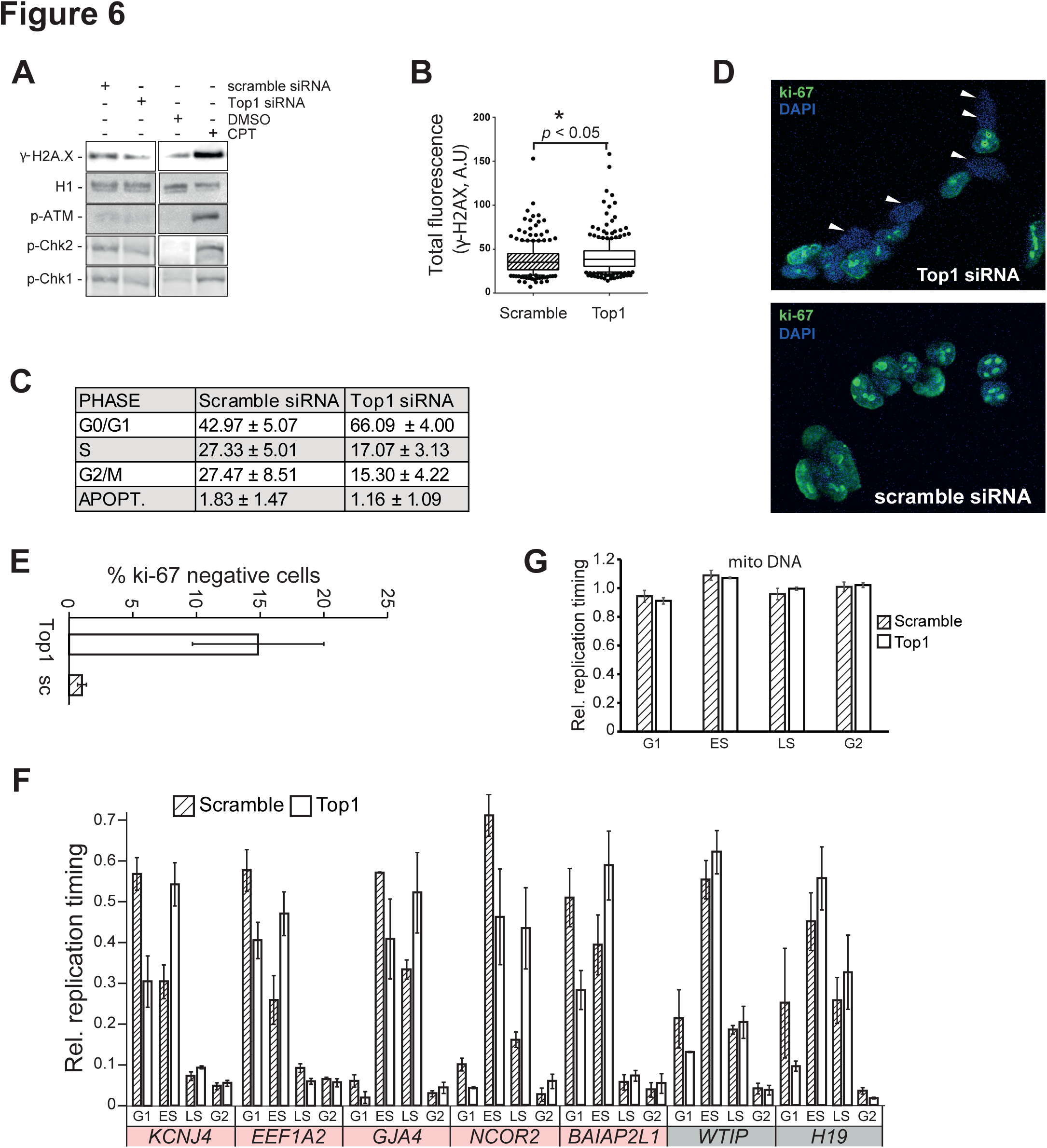
Top1 depletion triggers G0/G1 block and global replication timing delay. (A) Western blots showingthe cellular response to Top1 depletion and camptothecin treatment with respect to yH2AX and markers of DNA damage signaling. (B) Total ↃH2AX immunofluorescence signal for control and Top1-depleted cells (n >100 cells). (C) Cell cycle analysis for Top1-depleted cells and controls. Results are average of four experiments presented with standard deviation. (D) Representative images of ki-67 staining for Top1-depleted and control cells. Cells were counter-stained with DAPI. (E) Quantification of ki-67 staining (160 cells for each sample per experiment; two independent replicates). (F) Analysis of replication timing at a range of RLL loci in Top1-depleted cells and controls. The & of cells undergoing replication in each phase of the cell cycle was measured by the relative recovery of BrdU-labeled immunoprecipitated DNA across G1, early S (ES), late S (LS) and G2 phases. Error bars are SE of two replicates. Red and grey shading indicate genes with significant and non-significant replication timing delays, respectively. (G) Replication timing analysis of mitochondrial DNA.

The observation that Top1-depleted cells undergo a G1/S transition block, combined with the close association of RLL peaks with early, highly active, replication origins, suggested that Top1 depletion may impair the replication program. To test this, we analyzed the replication timing of multiple early-replicating RLL regions using BrdU incorporation to mark newly replicated strands, followed by immunoprecipitation and qPCR after cell sorting into G0/G1, early S, late S and G2/M phases. Top1-depleted cells showed delayed replication timing with a consistent switch from G1 to early S or from early S to late S phase for 5 out of 7 RLL loci (Fig. 6F). It should be noted however, that genes with no or mixed changes in R-loop distribution also showed a similar trend for 7 out of 8 loci tested (Supplemental Fig. S6B). A minority of RLG genes (3 out of 10) also showed a modest tendency towards replication delay (Supplemental Fig. S6C). This effect was specific for the nuclear genome, as mitochondrial DNA replication timing was not affected by Top1 knockdown (Fig. 6G). Thus, when cells are able to overcome the G1/G0 block and initiate S phase, they nonetheless show a trend towards replication delay in a way that appears influenced by, but not strictly dictated by, R-loop gains or losses.

## DISCUSSION

Multiple studies have identified Top1 as a factor that prevents R-loop formation since the enzyme relaxes negative supercoils during transcription [42, 43] thereby preventing an R-loop favorable underwound DNA state [7, 12]. To understand how Top1 modulates R-loop formation *in vivo*, we profiled these structures globally in Top1-depleted human cells. Surprisingly, Top1 depletion caused both increases and decreases of R-loop levels depending on the genomic context (Figs. 1 and Figs. 2).

Consistent with the expectation that Top1 prevents R-loops, we identified a clear class of genes that respond to Top1 depletion by gaining R-loops. These genes were long, highly transcribed, and located in gene-poor areas of the genome. These observations are consistent with prior studies and allow us to refine a model for Top1 activity during transcription elongation. Long genes were shown to be more sensitive to Top1 poisoning by Camptothecin [44] or to Top1 depletion in mouse and human neurons [45]. These studies are in agreement with our observations that RLG peaks arise co-transcriptionally on long and highly expressed genes, where they principally match to gene bodies (Figs. 1-3). Our work also shows that RLG genes preferentially undergo promoter stalling (Fig. 2) which is in agreement with a recent study showing that Top1 becomes physically associated with the RNAP complex and catalytically activated upon release of the transcription machinery into elongation from a promoter-proximal paused state [32]. Our work supports the view that Top1 facilitates transcription elongation and precisely defines the class of genes that are most dependent on this enzyme: Top1 efficiently prevents co-transcriptional R-loops specifically for long gene units with high transcription levels.

Interestingly, the generation of topological stress during transcription requires that the DNA fiber is placed under some physical constraints so as to prevent spontaneous dissipation of supercoils [1]. Our study reveals that in the case of R-loop stabilization through RLG genes, proximity to H3K9me3-marked chromatin and lamin-associated domains may represent the main source of such physical constraints. A subset of RLG genes were located in close proximity to LADs and faced away from the LAD boundary, suggesting that LADs might physically trap supercoils, causing an increase in negative supercoil density behind the transcribing RNAP in the absence of Top1. Indeed, the strength of R-loop gains clearly correlated with the proximity to LADs (Fig. 4). We therefore suggest that the association of genes to the nuclear envelope sensitizes them to topological disruptions. In addition to LADs, we identify heterochromatic H3K9me3-marked patches as a second important distinguishing feature of RLG genes. These patches often were in close proximity to RLG peaks and the strength of RLGs was inversely correlated to their distance from H3K9me3 peaks. We suggest that H3K9me3-marked heterochromatic patches might prevent dissipation of torsional tension because of their closed chromatin nature. Additionally, H3K9me3 was shown to mediate perinuclear anchoring, which could further prevent supercoil dissipation [46]. Altogether, our data reveals that long, highly expressed genes in proximity to LADs or H3K9me3 patches are important reservoirs of R-loops that require proper topological control by Top1. Top1 depletion may be less critical for genes without such topological constraints where activity of Top2 isoforms may be sufficient to substitute for Top1 absence.

Given the association between R-loops and RNAP pausing [4] as well as DNA breakage [3], we speculate that R-loop suppression by Top1 plays an important role in ensuring proper gene expression and genome stability. We note that, in contrast to other studies [14, 47], we did not detect telltale signs of genomic instability (Fig. 6). Importantly, these studies used cell lines in which Top1 was stably knocked down and that were forced to undergo replication by passaging and selection. By contrast, our study only involved transient Top1 knockdown and caused a strong G0/G1 cell cycle block (Fig. 6; see below). Given that passage through S phase is required for R-loop-induced DNA breakage and instability phenotypes [2, 14], the reduction of cells in S phase likely counteracted the accumulation of DNA damage in our cell model. We speculate, however, that RLG genes may represent a source of genomic instability once cells are able to replicate. We also note that Top1 depletion in our system did not result in major changes in gene expression (Supplemental Fig. S2D) or notable accumulation of RNAP at sites of RLG (data not shown). This indicates that, while transient Top1 depletion caused R-loop accumulation in RLG genes, gene expression still proceeded mostly unchanged. It is possible that the loss of Top1 activity, particularly in removing positive supercoils that might hinder RNAP progression, was compensated by the redundant activity of Top2. Recent evidence indeed shows that genes with high transcriptional outputs require Top2 activity to properly handle the resulting torsional stress [42]. Thus, unlike widely held views, Top1 depletion does not result in a global R-loop increase but rather affects a specific subset of genes. This study identifies RLG genes as uniquely Top1-responsive and reveals the molecular features that render these genes dependent on Top1 for R-loop control.

Unexpectedly, Top1 depletion also led to R-loop losses over a class of genes entirely distinct from RLG genes. RLL genes were of average length, resided in gene-rich neighborhoods, and were moderately expressed (Figs. 2, 4). The most striking feature of RLL peaks was their tendency to co-localize with replication initiation regions as defined either by SNS-seq or OK-seq (Fig. 5). This co-localization was underscored by the fact that RLL loci showed higher SNS-seq signal than matched or RLG loci. Initiation regions highlighted by their RLL overlap replicated early (predominantly G1), earlier than other Top1-invariant R-loop forming loci matched for gene expression and gene densities. Studies of replication origins in mammalian systems indicate that early origins are characterized by marks of open chromatin and by transcription [48-50]. Fittingly, co-transcriptional R-loops preferentially associate with increased DNase accessibility, histone H3 acetylation, and histone H3 lysine 4 methylation [4, 22]. Top1-responsive RLL peaks further include a significant association with the H3K27me3 Polycomb mark and components of the PRC complexes (Fig. 5A). Interestingly, a subset of early, highly efficient replication origins was previously associated with a very similar chromatin pattern [40, 49]. Thus RLL peaks correspond to Top1-responsive R-loop forming loci that are enriched over a subset of early active replication origins and preferentially carry chromatin marks previously defined for these loci.

Interestingly, a primary cellular response to Top1 depletion is the accumulation of cells in G1/G0 and a delay in replication timing at certain genomic loci. One possible mechanism to account for this observation is if Top1 and R-loops participate in origin function. Top1 is known to bind to replication origin sequences [39, 51] as part of the replication progression complex (RPC) which comprises the MCM and GINS proteins [52]. In the SV40 system, almost all RPC components are individually dispensable for activation of SV40 origin in crude extracts, except for Top1 and its interaction with the T antigen for the priming of viral replication [53, 54]. Top1 is therefore a part of the basal complex responsible for origin activation and nascent fork formation. Furthermore, Top1 DNA cleavage sites have been mapped at the lamin B2 origin and Top1 inhibition by low camptothecin concentrations abolished origin firing, suggesting that Top1 and DNA topology play a key role in this process [51]. A plausible hypothesis is that the catalytic activity of Top1 is necessary at replication origins to remove the positive, but not negative, supercoils generated by the unwinding of DNA mediated by MCM helicases [55], leaving the DNA template more negatively supercoiled and thus favoring DNA strand separation. If so, the absence of Top1 will cause the inefficient removal of positive supercoils which in turns will disfavor R-loop formation and cause the appearance of RLL loci. Our work therefore highlights that Top1 may play an important role in controlling replication origin function in human cells at a subset of early origins. It nonetheless remains possible that Top1 depletion may affect replication timing and cell cycle progression in an indirect and more complex manner; further investigations will be necessary to fully define the molecular mechanisms linking Top1 and replication origin activity.

In addition, our work also raises the possibility that R-loop formation may be linked to replication origin function in human cells. The notion that R-loops may contribute to origin function is supported by a wide array of observations. As mentioned above, R-loop forming regions associate with chromatin signatures that are typical of replication origins. R-loops and origins both show hotspots of distribution at gene ends [41, 56]. CpG island promoters in particular, are R-loop and origin hotspots [5, 50, 57-61], and associate with conserved patterns of GC skew [56, 62], a sequence characteristic that intrinsically favors the formation of G-rich signatures often referred to as origin G-rich repeated elements [63]. Such G-rich motifs have the potential to form G quadruplex structures that have been implicated as determinants of origin positioning and efficiency [63, 64]. While it is unclear if G quadruplex can spontaneously nucleate in the context of double-stranded DNA, it is reasonable to propose that R-loop structures can favor G4 formation on the looped out single-strand [65]. Interestingly, the ORC1 subunit of the origin Recognition Complex was shown to bind G4-preferrable ssDNA [66], thereby suggesting that R-loop formation may favor origin licensing. Several historical precedents further underscore the connections between R-loops and origins. In the T4 bacteriophage and in ColEI-replicons in *E. coli*, R-loops function as replication origins [25-27, 67]. In *E. coli*, recombination-mediated R-loops in RNase H-deficient strains support an OriC-independent mode of replication [68, 69]. Increased R-loop formation in RNase H-deficient yeast strains subjected to Top1 inhibition led to origin-independent DNA replication initiation in the rDNA [28]. Finally, the mitochondrial genome is thought to initiate DNA replication priming through R-loop intermediates [70-72] and a recent study showed that replication origins are specified in an R-loop dependent manner at murine class switch immunoglobulin regions [73]. Thus, as judged from location overlaps, chromatin features, and functional associations, our work is consistent with an intimate connection between R-loop formation and replication origin specification [74]. Future work will be necessary to delineate the detailed mechanistic connections that link transcription, R-loop formation, topoisomerase activity, and replication initiation. Altogether, our work establishes that Top1 functions at the interface between DNA replication and transcription and regulates R-loop formation in a context-dependent manner.

## METHODS

### Cell Lines and Drugs

HEK293 cells (ATCC) were maintained in DMEM (Thermofisher) supplemented with 10% FBS in a humidified incubator at 5% of CO_2_. Camptothecin (Sigma Aldrich) was dissolved in DMSO at 10 mM concentration, stored in aliquots at −20°, and used as a 1,000x stock during 1 hour treatments.

### Top1 Knockdown

HEK293 cells were counted and seeded at 150,000 cells per 35 mm dish. 24 hours after seeding, cells were reverse transfected using RNAimax transfection reagent (Thermofisher) and with 10 nM of Top1 specific siRNA (Thermofisher) targeting exon 16 (siRNA #1) and exon 15 (siRNA #2) of the Top1 transcript, or with a negative control RNA (scramble). 48 hours after the first transfection, one fifth of the cells were transfected again in a similar manner. Cells were harvested 72 hours after the second transfection for all subsequent analysis.

### Dot Blot Analysis

Genomic DNA was extracted according to DRIP protocol and digested with restriction enzyme cocktail mix. Serial dilutions starting from 7.5 micrograms of DNA were spotted on a nitrocellulose membrane and crosslinked with UV light (120 mJ/cm^2^). Membrane was blocked with PBS-Tween (0.1%) and 3% BSA for 30 min and then incubated with S9.6 antibody diluted to 1 μg/ml in PBS-Tween (0.1%), 3% BSA. After washing, membrane was incubated with HRP-conjugated or Alexa-fluor 488 anti-mouse secondary antibodies, further washed and developed with ECL techniques or directly in fluorescence scanning. In case of treatment with RNase H genomic DNA was pre-incubated with 10 U of enzyme for two hours at 37°C.

### Western Blot

Western blot analysis was performed according to standard procedures. Membranes were incubated with the following antibodies: anti Top1 (c15, sc5342), anti beta-actin (I-19, sc1616), anti p-ATM (10H11.E12, sc47739), anti histone H1 (AE-4, sc8030) from Santa Cruz Biotechnology. Anti Phospho-H2AX antibody (ser139, JBW301) was from Millipore. Anti Phospho-ChK1 (Ser345, 133D3) and anti Phospho-Chk2 (Thr68, C13C1) were from Cell Signaling.

### DRIP and DRIPc-seq

DRIPc-seq was performed as previously described [4]. Briefly, DRIP immunoprecipitates obtained from 40 micrograms of digested genomic DNA were collected and treated with DNase I (Fermentas). The resulting RNA strands were purified and reverse-transcribed using the iScript kit (Bio-Rad). Second strand synthesis was performed using dUTP instead of dTTP. Ligation of Illumina Truseq adapters was performed according to manufacturer’s instructions and a UDG glycosylase treatment was introduced before library amplification to permit strand-specific R-loop detection. In case of treatment with RNase H or RNase A, digested genomic DNA was pre-treated with 10 units of RNase H or 10 μg/ml of RNase A for two hours at 37°C before DRIP.

### RNA Pol II ChIP-seq and total RNA-seq

RNA Pol II ChIP was performed as previously described [75]. Immunoprecipitated DNA was purified and used to construct Illumina NGS libraries according to manufacturer procedures. Total RNA-seq was performed after ribosomal RNA depletion using an Illumina Truseq RNA-seq kit according to the manufacturer’s instructions.

### DRIPc-seq, RNA-seq, and RNA Pol II ChIP-seq Mapping and Peak Calling

Sequenced single-end reads were subjected to standard quality control pipeline using fastq-mcf software and mapped using Tophat2 for RNA-seq and Bowtie2 for the rest with default parameters. Sequencing read depths were normalized by number of mapped reads between samples, and only uniquely mapped reads were considered. High copy-number or contamination-prone regions such as rDNA, mitochondria, centromere, and ENCODE blacklisted regions were excluded. DRIPc-seq peak calling was performed using a previously developed Hidden Markov Model [4] modified to enable higher sensitivity in particular when dealing with lower and trailing signal (see https://github.com/srhartono/highsenshmm.) This method was about 2.5-fold more sensitive, generating about 200,000 peaks of signal covering about 500 MB of genomic space in each replicate. For analysis, all DRIPc-seq peaks present in at least one sample were considered and regions showing significant differences in signal between Top1-depleted and control cells were identified using DESeq2 using significance thresholds of an adjusted p-value < 0.1 and signal fold-change higher than 1.25x or lower than 0.8x (using a more stringent adjusted p-value < 0.05 did not affect our conclusions; data not shown). Genes shorter than 5 kb were eliminated from this analysis.

### Overlap Analysis with Other Datasets

Datasets for lamin, chromatin marks, ChromHMM states, SNS-seq and OK-seq replication origins or zones were downloaded from published sources (Supplemental Table 2). The RNAP pausing state of each gene was categorized as in [76] using RNA Pol II ChIP-seq datasets generated from control HEK293 cells. The enrichment or depletion of RLL and RLG peaks over chromatin features of interest was first measured in terms of peak overlap. For this, we determined the overlap of RLL and RLG peaks over chromatin peaks of interest and then calculated the peak overlap for control peaks. These control peaks were stringently selected following an earlier strategy [4]. In brief, these peaks belonged to expression-and length-matched R-loop forming genes that were not affected by Top1 depletion (no and mixed changes in Fig. 2A). In all cases, these peaks were matched to a similar-sized R-loop peak on the matched gene. In the case of promoters and terminators, the precise position of the initial and shuffled peaks was maintained. Each initial RLL or RLG peak was independently matched multiple times to avoid outliers. We next determined the ratio of overlaps between RLL or RLG peaks and control peaks and expressed this ratio as a heatmap. The absolute overlap of chromatin features with RLL and RLG peaks is indicated by stars, as shown in Figs. 4 and 5. SNS-seq origin peaks were from [40] for human K562 and HeLa cells. OK-seq data was downloaded as RFD values from [41] for human HeLa cells. The RFD signal was processed using an HMM model configured as described by Petryk et al. (2016) to call replication initiation zones. Overlap between SNS-seq origins and OK-seq initiation zones was measured relative to stringent matched controls. Given that R-loop mapping was performed in HEK293 cells, it is likely that the overlap with replication initiation regions was under-estimated.

### Immunofluorescence

48 hours after second round of transfection with siRNA oligonucleotides, cells were detached and seeded at 300,000 cells per 35 mm dish on a glass coverslip pre-treated with gelatin. 24 hours after seeding, cells were methanol fixed and treated with acetone. Blocking and ki-67 (Abcam, ab15580) or γH2AX antibody incubation were performed in 4x SSC and 3% BSA at 20°C for 30 min and 2 hours, respectively. Secondary antibody was anti-rabbit or anti-mouse Alexa-fluor 488. Nuclei were counter-stained with DAPI.

### Cell Cycle Analysis and Replication Timing

Cell cycle analysis and replication timing were performed as described previously [77]. Briefly, cells were pulse-labeled with BrdU (50 μM) for two hours. Cells were then harvested, fixed in 70% ethanol, and stored at −20°C. Before cell cycle analysis and sorting, cells were labeled with Propidium Iodide (50 μg/ml) and treated with RNase A (250 μg/ml). Cells were analyzed and sorted with a Biorad S3e cell sorter. After sorting, cells were lysed and genomic DNA was extracted. DNA was immunoprecipitated with anti BrdU antibody (B44, BD Biosciences, 347580), purified, and used as template in qPCR. To assess R-loop formation across the cell cycle, cells were first synchronized in G2 after thymidine block (24 hours) and released into nocodazole-containing media (12 hours). Cells were then allowed to cycle in fresh medium and harvested in G1 and mid-S of the following cycle for DRIP-qPCR analysis. Cytofluorometric analysis after propidium iodide staining confirmed that >85% of the cells were in the correct cell cycle phases.

## DECLARATIONS

### Data access

High-throughput sequencing information is available for download from NCBI GEO entry GSE102474.

## Authors contributions

S.M., F.C., and G.C. designed research. S.M. and L.S. performed experiments. S.R H. performed bioinformatics analysis. S.D.B. and A.C performed cell sorting analysis. S.M, S.R.H., F.C, G.C. analyzed the data and wrote the paper.

